# Intra-genome variability in the dinucleotide composition of SARS-CoV-2

**DOI:** 10.1101/2020.05.08.083816

**Authors:** Paul Digard, Hui Min Lee, Colin Sharp, Finn Grey, Eleanor Gaunt

## Abstract

CpG dinucleotides are under-represented in the genomes of single stranded RNA viruses, and coronaviruses, including SARS-CoV-2, are no exception to this. Artificial modification of CpG frequency is a valid approach for live attenuated vaccine development, and if this is to be applied to SARS-CoV-2, we must first understand the role CpG motifs play in regulating SARS-CoV-2 replication. Accordingly, the CpG composition of the newly emerged SARS-CoV-2 genome was characterised in the context of other coronaviruses. CpG suppression amongst coronaviruses does not significantly differ according to genera of virus, but does vary according to host species and primary replication site (a proxy for tissue tropism), supporting the hypothesis that viral CpG content may influence cross-species transmission. Although SARS-CoV-2 exhibits overall strong CpG suppression, this varies considerably across the genome, and the Envelope (E) open reading frame (ORF) and ORF10 demonstrate an absence of CpG suppression. While ORF10 is only present in the genomes of a subset of coronaviruses, E is essential for virus replication. Across the *Coronaviridae*, E genes display remarkably high variation in CpG composition, with those of SARS and SARS-CoV-2 having much higher CpG content than other coronaviruses isolated from humans. Phylogeny indicates that this is an ancestrally-derived trait reflecting their origin in bats, rather than something selected for after zoonotic transfer. Conservation of CpG motifs in these regions suggests that they have a functionality which over-rides the need to suppress CpG; an observation relevant to future strategies towards a rationally attenuated SARS-CoV-2 vaccine.

## Introduction

CpG dinucleotides are under-represented in the DNA genomes of vertebrates (Cooper and Krawczak 1989; Simmonds, et al. 2013). Cytosines in the CpG conformation may become methylated, and this methylation is used as a mechanism for transcriptional regulation (Medvedeva, et al. 2014). Methylated cytosines have a propensity to undergo spontaneous deamination (and so conversion to a thymine). Over evolutionary time, this has reduced the frequency of CpGs in vertebrate genomes (Cooper and Krawczak 1989). However, loss of CpGs in promoter regions would affect transcriptional regulation, and so CpGs are locally retained, resulting in functionally important ‘CpG islands’ found in around half of all vertebrate promoter regions (Deaton and Bird 2011).

Single strand RNA (ssRNA) viruses infecting vertebrate hosts reflect the CpG dinucleotide composition of their host in a type of mimicry (Simmonds, et al. 2013). It was hypothesised that this is because vertebrates have evolved a CpG sensor which flags transcripts with aberrant CpG frequencies (Atkinson, et al. 2014; Gaunt, et al. 2016). This idea was strengthened by the discovery that the cellular protein Zinc-finger Antiviral Protein (ZAP) binds CpG motifs on viral RNA and directs them for degradation (Takata, et al. 2017), and further supported by observations that CpGs can be synonymously introduced into a viral genome to the detriment of virus replication without negatively impacting transcriptional or translational efficiency (Tulloch, et al. 2014; Gaunt, et al. 2016). Current understanding is therefore that ssRNA viruses mimic the CpG composition of their host at least in part to subvert detection by ZAP. ssRNA viruses also under-represent the UpA dinucleotide, but to a far more modest extent (Simmonds, et al. 2013), and the reasons behind UpA suppression are less well understood. A consequence of dinucleotide bias is that certain codon pairs are under-represented (Tulloch, et al. 2014; Kunec and Osterrieder 2016) (so, for example, codon pairs of the conformation NNC-GNN are among the most rarely seen codon pairs in vertebrates (Tats, et al. 2008)). Whether the two phenomena of CpG suppression and codon pair bias (CPB) are discrete remains controversial (Futcher, et al. 2015; Kunec and Osterrieder 2016; Groenke, et al. 2020).

The *Coronaviridae* have a generally low genomic cytosine content (Berkhout and van Hemert 2015), but as with other ssRNA viruses, nonetheless still under-represent CpG dinucleotides to a frequency below that predicted from individual base frequencies of cytosine and guanine (Woo, et al. 2007).

The Coronavirus family comprises four genera – the alpha, beta, gamma and delta-coronaviruses. Human-infecting coronaviruses (HCoVs) have been identified belonging to the alpha and beta genera (Hu, et al. 2015). Alphacoronaviruses infecting humans include HCoV-229E and the more recently discovered HCoV-NL63 (van der Hoek, et al. 2004). Betacoronaviruses include HCoV-OC43, HCoV-HKU1 (Woo, et al. 2005), severe acute respiratory syndrome (SARS)-CoV (Rota, et al. 2003), Middle East respiratory syndrome (MERS)-CoV (Zaki, et al. 2012) and the recently emerged SARS-CoV-2 (Lu, et al. 2020; Zhu, et al. 2020). Prior to the emergence of SARS-CoV-2, SARS-CoV had the strongest CpG suppression across human-infecting coronaviruses (Woo, et al. 2007). The reason(s) for this are uncertain, but loss of CpG from a virus genome upon zoonotic transfer into the human host has previously been reported for influenza A virus (Greenbaum, et al. 2008), potentially indicating an advantage of reduced CpG content for infection of the human respiratory tract. All human-infecting coronaviruses are thought to be derived from ancestral bat viruses, though intermediate hosts may have facilitated zoonotic passage in some cases (Banerjee, et al. 2019).

During replication, coronaviruses synthesise transcriptionally active negative sense sub-genomic RNAs which are of varying length. Sub-genomic RNAs are synthesised by the viral polymerase copying the genome up to a 5’ leader sequence (Liao and Lai 1994) which is repeated upstream of most open reading frames (ORFs) in the coronavirus genome (such repeats are referred to as transcription regulation sequences (TRSs)); this complementarity allows viral polymerase jumping from the 5’ leader sequence to directly upstream of ORFs preceded by a TRS (Sawicki and Sawicki 1998). The negative sense sub-genomic RNAs serve as efficient templates for production of mRNAs (Sawicki, et al. 2007). Generally, only the first ORF of a sub-genomic mRNA is translated (Perlman and Netland 2009), although leaky ribosomal scanning has been reported as a means for accessing alternative ORFs for several coronaviruses including SARS-CoV (Schaecher, et al. 2007).

SARS-CoV-2 was recently reported to have a CpG composition lower than other members of the betacoronavirus genus, comparable to certain canine alphacoronaviruses; an observation used to draw inferences over its origin and/or epizootic potential (Xia 2020). Here we show that coronaviruses have a broad range of CpG composition which is partially host and tissue tropism dependent, and that there is no difference in CpG content across coronavirus genera. There is however a striking disparity in CpG composition between SARS-CoV-2 ORFs, with the Envelope (E) protein ORF and ORF10 over-representing CpG dramatically. E ORF and ORF10 also have higher UpA dinucleotide composition and lower CPB scores than other ORFs. E ORF displays CpG suppression in all human-infecting viruses except SARS-CoV and SARS-CoV-2, suggesting a potential correlation between CpG presentation and disease severity in human-infecting coronaviruses.

## Results

### CpG suppression within coronavirus genomes varies between host species and tissue tropism but not between genera

The genomic CpG composition of all complete genome coronavirus sequences (*n* = 3407; downloaded and further processed as described in the methods section and **Fig 1**) were calculated using observed: expected (O:E) ratios, with any value below 1 indicating CpGs are under-represented relative to the genomic content of cytosine and guanine bases. A substantial range in GC content (from ∼ 0.32 – 0.47) was seen across the *Coronaviridae*, and as expected, all viruses exhibited some degree of CpG suppression, with CpG O:E ratios ranging from 0.37 to 0.74 (**Fig 2A**). To investigate the root of this variation, the coronavirus sequence dataset was refined to remove sequences with more than 90% nucleotide identity to reduce sampling biases (so, for example, SARS-CoV sequences of human origin were stripped from over 1000 representative sequences to just one). The CpG compositions of the remaining 215 sequences (**Table S1**) were compared between coronavirus genera (alpha, beta, gamma and delta). For the 215 representative sequences, a genus could be assigned for 203. No differences in CpG composition between coronavirus genera were apparent, although the gamma genus exhibited a tighter range (**Fig 2B**). Next, we examined whether differences in CpG composition between viruses isolated from different hosts explained the range in CpG composition across the *Coronaviridae*. For the 215 representative sequences, a host could be assigned to 210. Coronavirus sequences were divided into host groups, and groups with at least three divergent sequences were compared; this included bat, avian, camelid, canine, feline, human, mustelid, rodent, swine and ungulate viruses. Variation in CpG composition between coronaviruses detected in different host species was evident across groups (*p* = 0.0057) and between groups, with coronaviruses detected in canine and human species having lower CpG content and rodent and bat coronaviruses having the highest (**Fig. 2C**). Significant differences in CpG composition were detected between bat and canine (*p* = 0.0001), avian and rodent (*p* = 0.005), canine and mustelid (*p* = 0.011), canine and rodent (*p* < 0.0001), human and rodent (*p* = 0.002), and rodent and ungulate (*p* = 0.0026) viruses. All frequency ranges overlapped however, indicating viral CpG frequency alone seems to be a poor predictor of virus origin, contradicting the recent suggestion of a canine origin of SARS-CoV-2 (Xia 2020). Where sequences in a host group representative of both alpha and betacoronaviruses were available (which was the case for bat, camelid, canine, human, rodent and swine viruses), these sequences were split by genus and compared to determine whether coronavirus genera influenced coronavirus CpG frequencies in a host species-specific manner. By this method, the lack of difference in CpG composition of coronaviruses of different genera was maintained (**Fig. 2D**).

**Figure 1.**
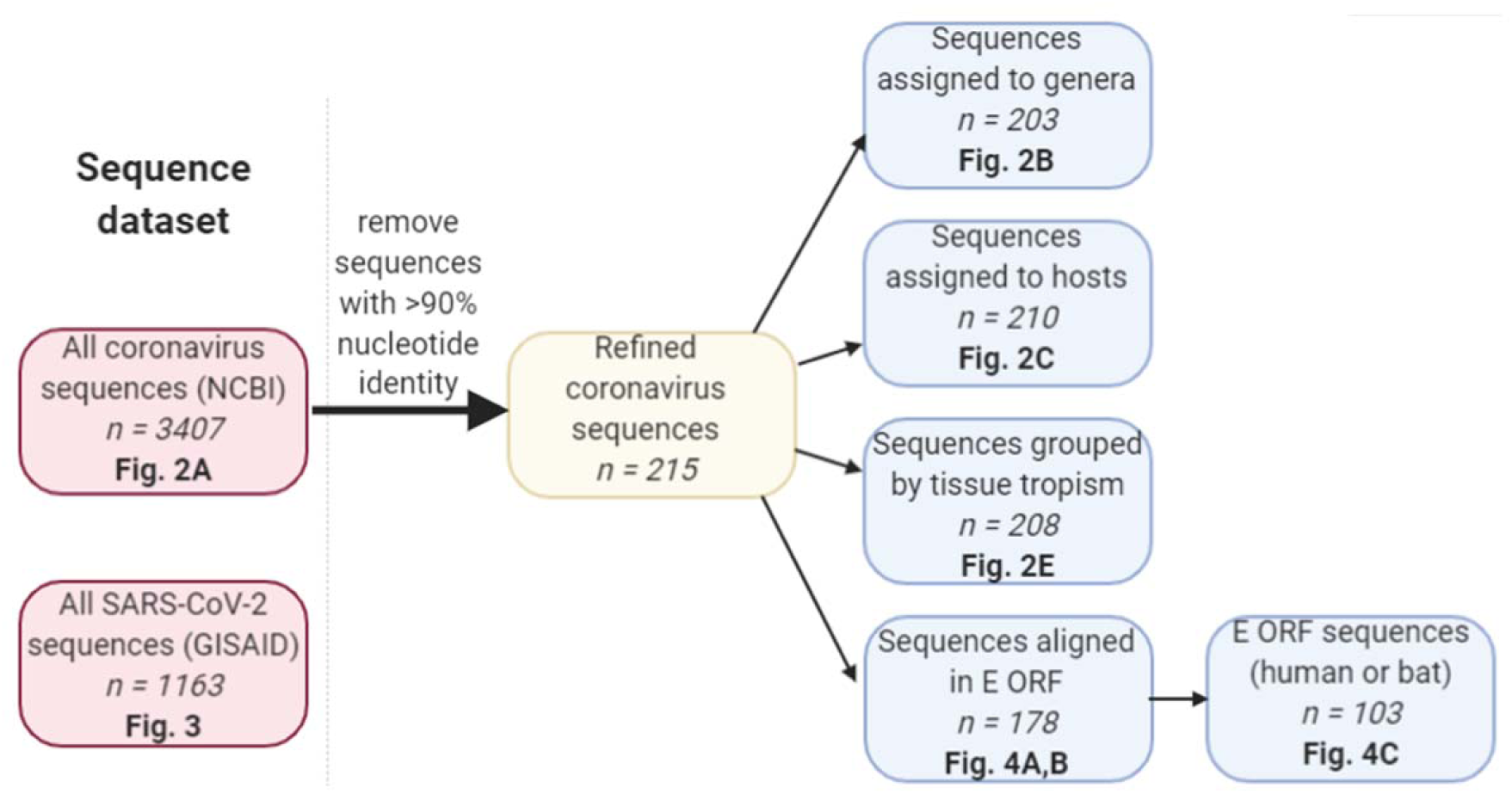
Workflow for sequence processing. Two sequence datasets were used for analysis; all coronavirus complete genome sequences available on NCBI, and SARS-CoV-2 complete genome sequences available on the GISAID platform (left hand pink shaded boxes). The coronavirus complete genome sequences were cleaned by removal of sequences with 90% nucleotide identity or greater to remove epidemiologic biases, leaving 215 complete genome sequences (central yellow shaded box). These were then categorised by genera, host, and tissue tropism. The subset of 215 sequences were also aligned over the E ORF and grouped by host (blue shaded boxes). Each box firstly describes each dataset used, the number of sequences in that dataset is then indicated in *italicized* font, and the figure to which the dataset corresponds is indicated in **bold** font.

**Figure 2.**
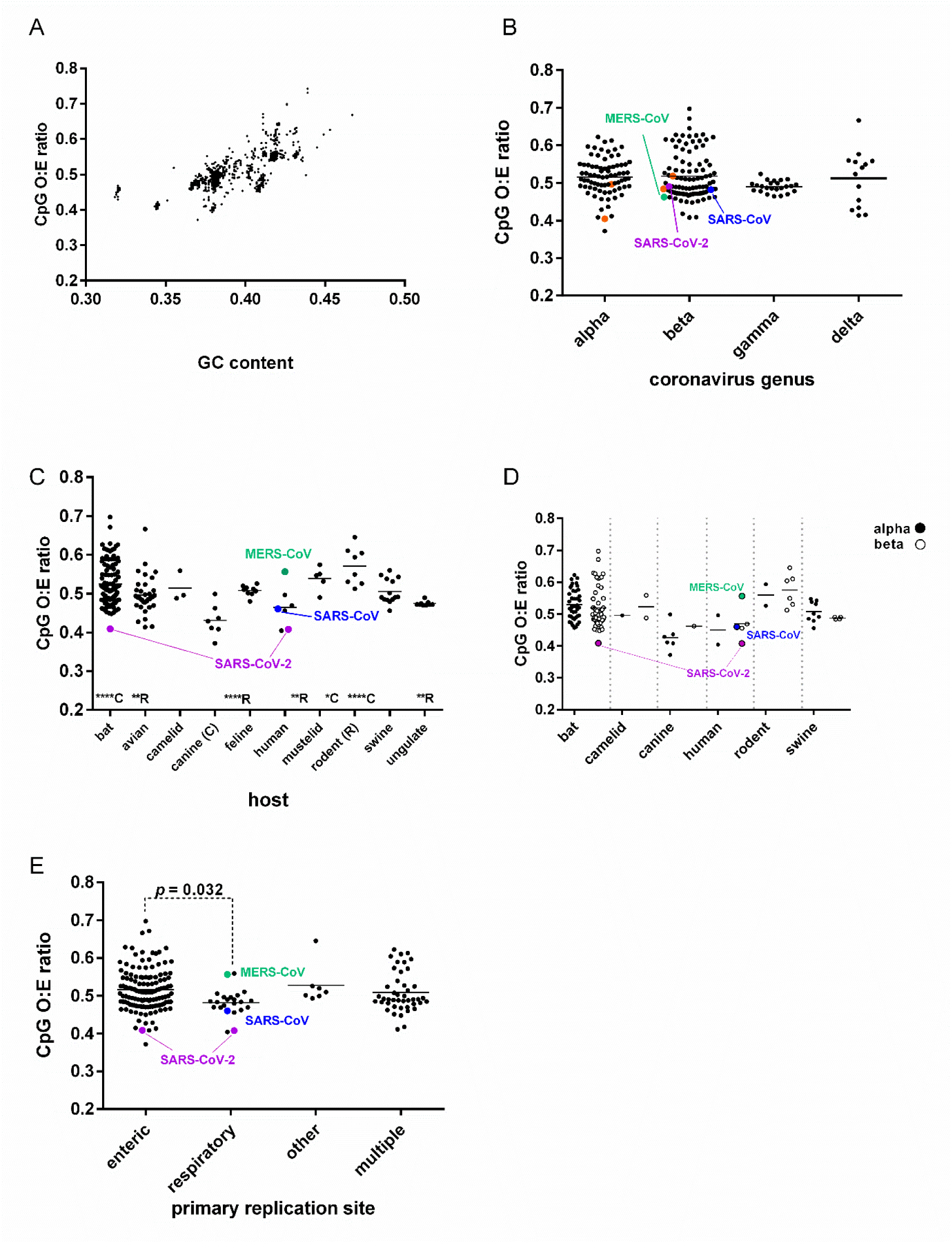
Comparison of the CpG ratios of complete genomes of coronaviruses. SARS-CoV is represented by a blue circle, SARS-CoV-2 by a purple circle and MERS-CoV by a green circle throughout. **A**. GC content versus CpG ratio for all complete genome sequences of coronaviruses downloaded from Genbank (3407 sequences). The sequence dataset in (A) was then stripped to include only one representative from sequences with less than 10% nucleotide diversity to overcome epidemiologic biases (215 representative sequences), which were analysed in the subsequent sub-figures. **B**. Coronavirus genus against genomic CpG content. Other human-infecting coronaviruses (HCoV-2292E, HCoV-NL63 (alphacoronaviruses) and HCoV-HKU1 and HCoV-OC43 (betacoronaviruses) are represented using orange circles. **C**. Vertebrate host of coronavirus against genomic CpG content. Statistically significant differences between CpG compositions of viruses from different hosts are indicated above the x axis line, with ‘C’ denoting a statistically significant difference from canine coronaviruses and ‘R’ denoting a statistically significant difference from rodent coronaviruses. Tukey’s multiple comparisons test was used to identify differences in CpG composition between viruses infecting different hosts. A *p* value < 0.05 is indicated with *, p < 0.01 = **, p < 0.001 = *** and p <0.0001 = ****. **D**. Vertebrate host of coronavirus, with further sub-division into coronavirus genus, against genomic CpG content. Alphacoronaviruses are denoted with filled circles and betacoronaviruses with open circles. **E**. Primary replication site against genomic CpG content by host. Tukey’s multiple comparisons test was used to identify differences in CpG composition between viruses infecting different tissues. For a full breakdown of how these were assigned, please refer to **Table S1**.

To test the hypothesis that coronavirus CpG content varies according to tissue tropism (Xia 2020), we classified the viruses according to their primary site of replication, where this was known or could be inferred from the sampling route. Samples were split into five categories – ‘respiratory’, ‘enteric’, ‘multiple’, ‘other’, or ‘unknown’. Altogether, 206 of the 215 sequences were classifiable (detailed in **Table S1**), with 9 sequences categorised as ‘unknown’ and excluded from further analyses. By this admittedly inexact approach, viruses infecting the respiratory tract had a significantly lower mean CpG composition than viruses with enteric tropism (*p* = 0.032; **Fig. 2E**). However, the spread of respiratory virus CpG frequencies was contained entirely within the range exhibited by enteric viruses. Furthermore, 124 sequences were assigned to the enteric group, and only 22 to the respiratory group. Of these 146 sequences, bat viruses accounted for 80, all of which were assigned to the enteric group (despite reasonable sampling of respiratory tract in bats) and this cohort of viruses maintained almost the full spread of CpG frequencies (**Fig. 2E, Table S1**). Thus, while coronavirus CpG frequency may show some correlation with replication site, the dataset available does not permit strong conclusions to be drawn or predictions about zoonotic potential to be made.

### Heterogeneities in the dinucleotide composition of SARS-CoV-2

By our methods for calculating CpG O:E ratios, SARS-CoV-2 has a genomic CpG ratio of 0.408 (representing the mean of 1163 complete genome sequences). This is similar to the value calculated previously for a much smaller sample (*n* = 5) of SARS-CoV-2 sequences (Xia 2020). As this previous study noted, this is at the bottom end of the range of genomic CpG O:E ratios for betacoronaviruses and for coronaviruses detected in humans (**Figs 2B, C and D**). However, as noted above, vertebrate DNA genomes contain localised islands of higher CpG content (Deaton and Bird 2011). To determine if similar heterogeneity in CpG frequency was evident in the SARS-CoV-2 genome, the composition of individual ORFs was examined. Overall, most ORFs had CpG O:E ratios which were comparable to the genomic CpG ratio. However, two ORFs in particular, E ORF and ORF10, had CpG ratios higher than 1, indicating an absence of CpG suppression in those regions (**Fig. 3A**). These two ORFs also did not suppress the UpA dinucleotide, in contrast with other SARS-CoV-2 ORFs (**Fig. 3B**).

**Figure 3.**
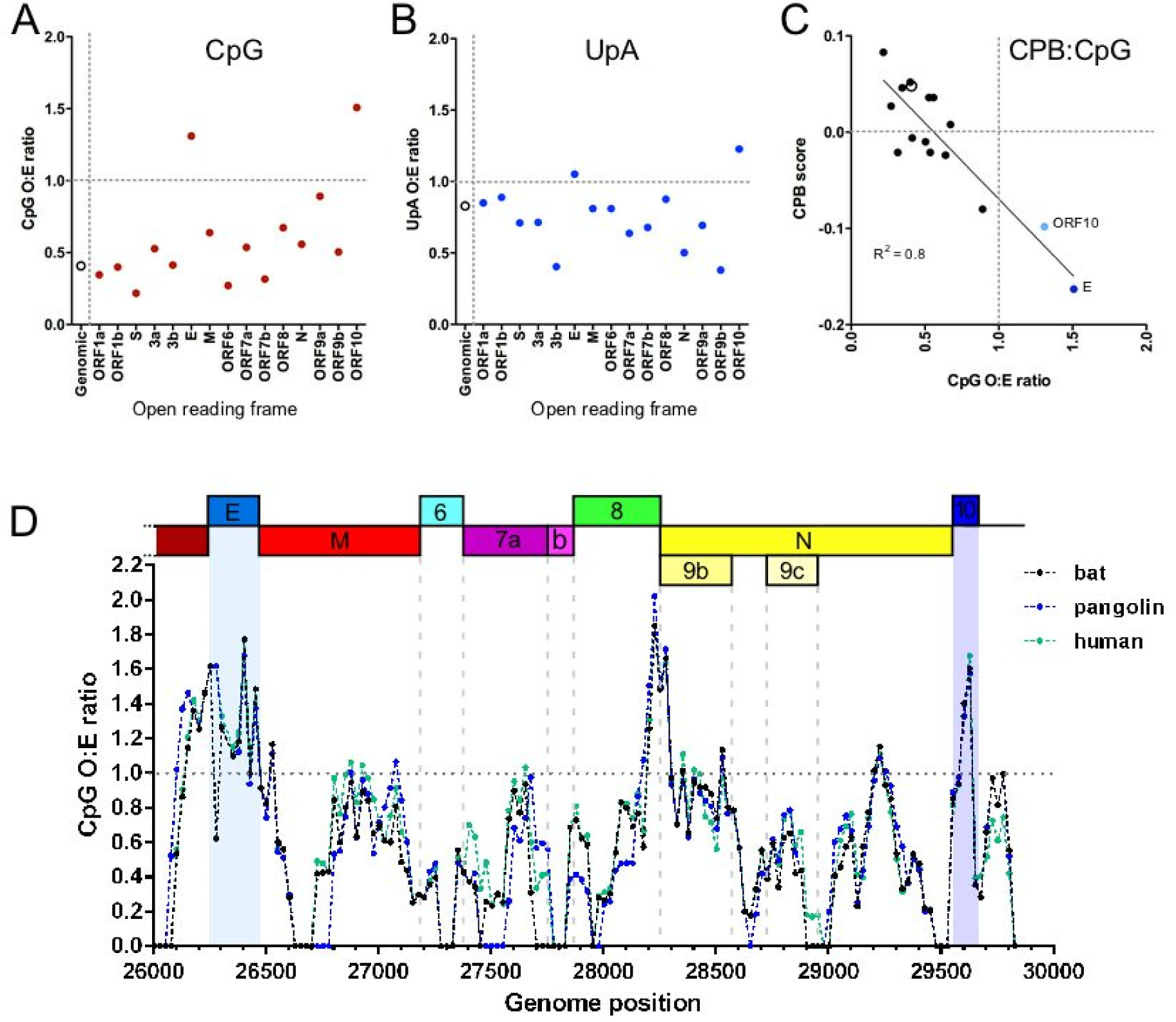
Heterogeneities in the dinucleotide composition of the SARS-CoV-2 genome. **A-C**. Comparison of the dinucleotide and coding compositions of SARS-CoV-2 open reading frames (ORFs) for **A**. CpG observed: expected (O:E) ratios, **B**. UpA O:E ratios and **C**. Codon pair bias (CPB) scores. Average scores across the genome are indicated using open circles. **D**. Sliding window analysis of CpG content of SARS-CoV-2 (green line) and closely related bat (black line) and pangolin (purple line) isolates. The CpG O:E ratio of the 3’ end of the genome was measured in 100 nucleotide windows in 25 nucleotide increments. The mean of 1163 complete genome sequences is presented for SARS-CoV-2.

Due to the difficulties in distinguishing between dinucleotide bias and CPB, CPB scores were also calculated for each ORF and plotted against CpG composition (**Fig. 3C**). CPB scores provide an indication of whether the codon pairs encoded in each ORF are congruous with usage in vertebrate genomes. A score below 0 indicates use of codon pairs that are disfavoured in host ORFs. An approximately linear relationship between CpG O:E ratio and CPB score for each SARS-CoV-2 ORF was apparent (R^2^ = 0.80). E ORF and ORF10 both had negative CPB scores, indicating that they use under-represented codon pairs and in keeping with the observation that both ORFs over-represent CpG and UpA dinucleotides.

To examine the precise location of the CpG hotspots, a sliding window analysis of CpG content across the 3’ end of the SARS-CoV-2 genome (averaged over 1163 complete genome sequences) as well as the closely related bat and pangolin sequences was performed. As expected, marked increases in CpG O:E ratio were observed concomitant with the genomic regions associated with E ORF and ORF10 (**Fig. 3D**). The E ORF and ORF10 regions associated with high CpG composition were maintained across the bat, pangolin and human sequences, indicating that since the bat sample was collected in 2013, the higher CpG frequency in this region has not been negatively selected. While the increase in CpG presentation was apparent across the entire E ORF, starting at the 3’ end of ORF3 and ending at the beginning of the M gene, the CpG spike in ORF10 was more narrowly associated with the putative coding region. Additionally, a CpG spike between the 3’-end of ORF8 and the 5’-end of the N gene was evident. The 5’-end of the N ORF also contains the overlapping ORF9b gene, which when considered alone, has a CpG O:E ratio approaching 1 (**Fig. 3A**), and is the ORF with the third-highest CpG O:E ratio after E ORF and ORF10. The usual coding plasticity afforded to nucleotides in the third position of a codon is nullified when overlapping reading frames are present, and so the CpG spike at this gene boundary is not surprising. Thus, although the SARS-CoV-2 genome exhibits high CpG suppression overall, there are local heterogeneities associated with individual ORFs, most notably E.

### On the origins of the high CpG content of E ORF of SARS-CoV-2

To determine whether the high CpG content of E ORF is evolutionarily conserved (ORF10 is poorly conserved and only encoded by a subset of SARS-like coronaviruses, so it was not analysed), attempts to identify the E ORF by nucleotide alignment for the set of 215 coronavirus sequences was undertaken, compared with E ORFs already annotated in NCBI. Of the 215 sequences, E ORF was identifiable in 178, with the remaining sequences too divergent to be confident of gene assignment. CpG composition for E ORF for the 178 sequences was measured and plotted according to host (**Fig. 4A**). A diverse distribution of CpG content was evident in viruses from every host group except ungulates, with bats in particular displaying a notable range from total suppression to overrepresentation. Otherwise, most viruses from most species still maintained some level of CpG suppression in E ORF. Overall, there was a significant difference between the mean CpG O:E ratio across host groups by 1-way ANOVA (p < 0.0001), but no significant differences were identified in pairwise comparisons of viral CpG compositions between hosts. The exceptions with high CpG O:E ratios in E ORF were four avian coronaviruses and notably, SARS-CoV and SARS-CoV-2. In contrast, other human-infecting coronaviruses (HCoV-229E, HCoV-HKU1, HCoV-NL63 and HCoV-OC43) all strongly under-represented CpG in E ORF, while MERS-CoV E ORF had an intermediate CpG O:E ratio of 0.6.To confirm E ORF over-represented CpG relative to the rest of the genome in SARS-CoV and SARS-CoV-2, ratios for E ORF: genomic CpG O:E were calculated (**Fig. 4B**). In non-bat non-avian host genomes, E ORF usually displayed CpG suppression in line with or stronger than that seen at the genome level, whereas SARS-CoV and SARS-CoV-2 starkly contrasted with this, displaying far less CpG suppression in this region.

**Figure 4.**
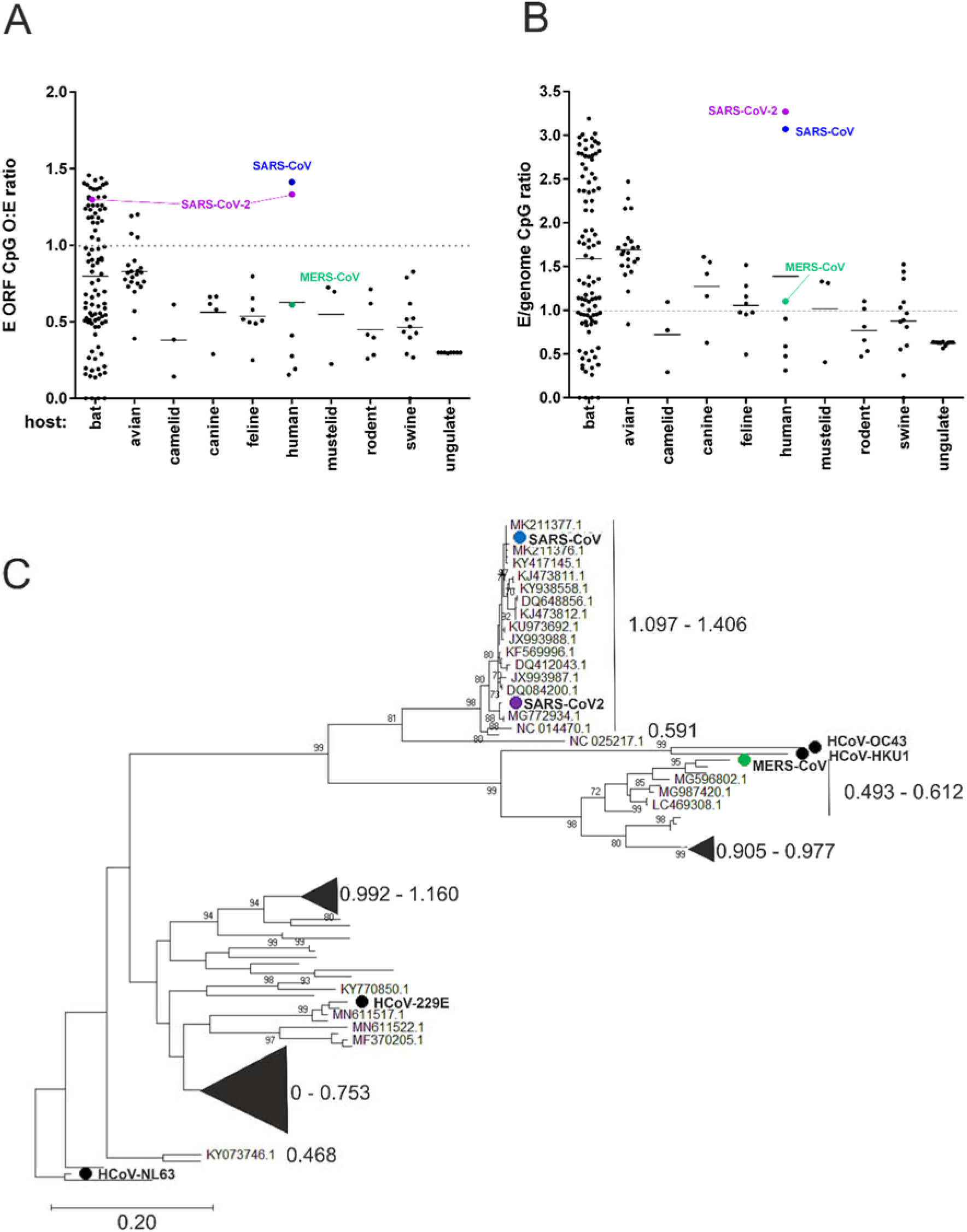
Evolutionary conservation of E ORF CpG content. MERS-CoV (green circle), SARS-CoV (blue circle) and SARS-CoV-2 (green circle) are indicated in both panels. **A**. Virus-dependent variation in E ORF CpG frequency. CpG O:E ratios for E ORF for 178 coronavirus E ORFs are plotted by host. **B**. CpG O:E ratios for E ORF were divided by the genomic CpG O:E ratio for 178 coronavirus sequences and grouped by host. **C**. Phylogenetic reconstruction of E ORF of human and bat coronaviruses. Maximum composite likelihood tree (100 bootstraps) representing the seven human-infecting coronaviruses (HCoV-229E, HCoV-HKU1, HCoV-NL63, HCoV-OC43 (all black circles) and 96 bat coronaviruses for which E ORF could be identified by alignment with the human coronaviruses. CpG O:E ratios for the E gene are indicated by large font numbers, and the sequences to which they relate are either bracketed or represented by triangles scaled to indicate the number of sequences they represent.

To investigate the evolutionary history of E ORF CpG composition in the human-infecting coronaviruses, a phylogenetic reconstruction of all 7 human coronavirus and 96 bat coronavirus E genes was performed to determine whether CpG ratios in this region were ancestrally derived. As expected (Cotten, et al. 2013; Lu, et al. 2020), the human viruses were interspersed among the bat viruses, reflective of their independent emergence events (**Fig. 4C**). The CpG compositions of the human coronavirus E ORFs, although diverse, were similar to the CpG compositions of their nearest bat relatives, demonstrating that CpG composition in E ORF is an ancestrally derived trait selected prior to emergence in the human population.

## Discussion

We have examined the CpG O:E ratios of all the currently available complete genome sequences of coronaviruses and uncovered a noteworthy diversity. Generally, the CpG O:E ratio of coronavirus genomes from a single host species varied considerably. For bats, which serve as a coronavirus reservoir (Banerjee, et al. 2019) and which had the largest number of representative sequences, the CpG O:E range was from 0.41 to 0.70, demonstrating the genome plasticity of coronaviruses and indicating that their evolution is not overtly restricted by a requirement to minimise CpG composition in the natural reservoir. The antiviral CpG-detector protein, ZAP (Takata, et al. 2017), has been identified as a target for several viral proteins including the 3C protease of enterovirus 71 (Xie, et al. 2018) and NS1 of influenza A virus (Tang, et al. 2017) – two viruses with overall low CpG content (Atkinson, et al. 2014; Gaunt, et al. 2016). This highlights the importance of CpG as a pathogen-associated molecular pattern (PAMP), and so this diversity in CpG expression within the *Coronaviridae* is striking. If coronaviruses also produce a protein with anti-ZAP activity, it is possible that this has variable efficacy between strains, explaining the ability of coronaviruses to fluctuate CpG composition considerably. Alternatively (or in addition), this may be host driven; we show that average CpG suppression varies with host species (**Fig 2C**) and, as previously suggested (Xia 2020), this may be linked with ZAP expression levels. We have demonstrated that CpG variation is not related to viral taxonomic grouping (**Fig. 2B**) but we did find an association between viral CpG composition and primary replication site, with respiratory coronaviruses having a lower CpG composition than enteric ones (**Fig. 2E**). This is the opposite of what has been previously suggested (Xia 2020), though this proposal was not supported by any comprehensive investigation.

Nevertheless, our meta-analysis was subject to the sampling preferences of many labs who have performed surveillance for coronaviruses, and many of the tissue tropism assignments we made have not been verified by experimental infections. Another limitation of this analysis is that only sequences of greater than 10% divergence were included. Tissue tropism can be defined by much smaller differences; for example, a deletion in the spike protein of transmissible gastroenteritis virus (a porcine coronavirus) altered the tropism of the virus from enteric to respiratory, while nucleotide identity was preserved at 96% (Cox, et al. 1990; Rasschaert, et al. 1990). Further study on tissue tropisms of coronaviruses, as well as tissue expression profiles and antiviral activities of ZAP are needed to validate these analyses.

Loss of CpG motifs during adaptation to the human host has been previously described for influenza A virus (Greenbaum, et al. 2008), highlighting the importance of CpG composition for host adaptation. For SARS-CoV-2, we determined a genomic CpG O:E ratio of 0.408, which is similar to the human genome CpG O:E ratio of 0.2-0.4 (McClelland and Ivarie 1982; Sved and Bird 1990; Tomso and Bell 2003). Mimicry of the CpG composition of the host by ssRNA viruses is considered a mechanism to subvert detection by the innate immune response (Simmonds, et al. 2013; Takata, et al. 2017) and speculatively this may indicate that SARS-CoV-2 was genetically predisposed to make a host switch into humans. Similarly, the genomic CPB score of 0.048 indicates that SARS-CoV-2 uses codon pairs which are preferentially utilised in the human ORFeome, which may mean that the virus was well suited for translational efficiency in humans at its time of emergence.

In coding regions which do not have overlapping ORFs, there is no requirement at the coding level for CpG motifs to be retained (Kanaya, et al. 2001). E ORF and ORF10 are not known to be in overlapping reading frames; conversely, ORF9b overlaps with the ORF for nucleocapsid (N). Some CpG retention in this region is therefore inevitable and may explain the high CpG composition of ORF9b. This nevertheless leaves open the question of why CpG motifs are retained in the E ORF and ORF10 regions (if this is not an ancestrally derived evolutionary hangover; as CpGs have not been lost from these regions between 2013 and now (**Fig. 3D**), this seems unlikely). CpG motifs may serve various non-exclusive purposes, including providing secondary structure (Rima and McFerran 1997), intentionally stimulating ZAP activity (by analogy with multiple viruses intentionally triggering NF-kB (Hiscott, et al. 2001)), or providing m5c methylation sites (Squires, et al. 2012; Khoddami and Cairns 2013; Dev, et al. 2017).

It is also possible that CpG enrichment serves as a strategy for regulating translation. Conceivably, the high CpG content at the 5’ end of the E ORF transcript destines this for degradation via ZAP or CBP-associated mechanisms (Guo, et al. 2007; Groenke, et al. 2020) more rapidly than other viral transcripts. This could be intentional, or an evolutionarily accepted trade-off to preserve a higher importance role for CpGs. Alternatively, E ORF and ORF10 proteins may only be required late during infection (parallels with which can be drawn from the differential temporal expression and translational efficiencies of transcripts of the coronavirus mouse hepatitis virus strain A59 (Irigoyen, et al. 2016)), by which time an as-yet unidentified inhibitor of ZAP (or other CpG/CBP sensor(s)) may render CpG suppression unnecessary, as suggested for human cytomegalovirus (Lin, et al. 2020).

ORF9b and ORF10 do not have their own TRSs and so whether or how these open reading frames are accessed is currently controversial; nevertheless, peptides from both have been identified by mass spectrometry from SARS-CoV-2 infected cells (Davidson, et al. 2020). The ORF9b AUG transcription initiation site, which has a strong Kozak context (Kozak 1986), is the first AUG after and 10 nucleotides downstream of the initiation site for N ORF (which displays moderate Kozak context). It is therefore credible to think that ORF9b is accessed via leaky ribosomal scanning -a well characterised method for accessing alternative ORFs used by coronaviruses and other viruses (Lin and Lo 1992; Chenik, et al. 1995; Schneider, et al. 1997; Senanayake and Brian 1997; O’Connor and Brian 2000; Ryabova, et al. 2006; Firth and Atkins 2010; Wise, et al. 2011; Irigoyen, et al. 2016). There is a lack of evidence that ORF10 is accessed via production of its own subgenomic RNA (Kim, et al. 2020); possibly, this ORF is accessed via leaky scanning from the leader immediately preceding the N ORF. However, visual inspection of the SARS-CoV-2 genome indicated that the AUG encoding ORF10 is 24 AUGs downstream from the one initiating N ORF, making this hypothesis speculative at best. Whether the anomalous CpG composition of ORF10 is somehow involved in priming its transcription remains to be determined.

The transcript encoding E ORF incorporates an additional ∼3.4kb of RNA and ORF10, if accessed from the transcript produced from the TRS upstream of N ORF, is present on a transcript of approximately 1.6kb in length. Whether the described CpG enriched regions are relevant as PAMPs in these contexts is currently unclear from what is known about ZAP recognition of CpG motifs. It is also worth noting that the body TRS sequence ahead of the E gene is relatively weak in SARS-CoV-2, as it is in SARS-CoV (Marra, et al. 2003), suggesting that this subgenomic mRNA may be of relatively low abundance. Of the SARS-CoV-2 transcripts which use a canonical TRS for synthesis, the donor site upstream of E ranked seventh when comparing sequencing read frequency across this site (behind reads spanning the TRS sites upstream of N, spike, ORF7a, ORF7b, ORF3a, ORF8 and M ORF respectively) in Vero cells infected at a low MOI for 24 hours, indicating that E ORF is of lower abundance than most other transcripts (Kim, et al. 2020). It is therefore possible that E ORF is of sufficiently low abundance for a high CpG frequency to be physiologically inconsequential. Similar logic can be applied to ORF10, which is just 117 nucleotides in length.

Synonymous addition of CpGs into a virus genome has been suggested as a potential novel approach to vaccine development by us and others (Burns, et al. 2009; Atkinson, et al. 2014; Gaunt, et al. 2016; Moratorio, et al. 2017). Here we explore the evolutionary space occupied by coronaviruses in the context of their CpG composition and find that SARS-CoV-2 has a low CpG composition in comparison with other coronaviruses, but with CpG ‘hotspots’ in genomically disparate regions. This highlights the potential for large scale recoding of the SARS-CoV-2 genome by introduction of CpGs into multiple regions of the virus genome as a mechanism for generation of an attenuated live vaccine. Introduction of CpG into multiple sites could also be used to subvert the potential of the virus to revert to virulence through recombination. A challenge of live attenuated vaccine manufacture is to enable sufficient production of a vaccine virus that has a replication defect. Strategic introduction of CpGs into specific regions of the virus genome has the potential to negate a replication defect in ZAP knockout cells, if regions such as conserved secondary structures and overlapping reading frames are avoided (Ficarelli, et al. 2019; Odon, et al. 2019), thus allowing the generation of high titre replication-defective vaccine virus stocks.

## Materials and Methods

### Sequences

For a comparison of GC content versus CpG ratio, all SARS-CoV-2 complete genome sequences of high coverage (as defined on the GISAID website) were downloaded from GISAID (www.gisaid.org) on 26 March 2020 (1163 sequences in total) and aligned against the SARS-COV-2 reference sequence (Accession number NC_045512) using Simmonics software (Simmonds 2012) SSE v1.4 (pre-release download kindly provided by Prof. Peter Simmonds, Oxford University). All sequences represented human isolates except for one sequence of bat origin (hCoV-19/bat/Yunnan/RaTG13/2013; EPI_ISL_402131) and one sequence from a pangolin (hCoV-19/pangolin/Guangdong/1/2019; EPI_ISL_410721). All complete genome sequences of all coronaviruses were downloaded from NCBI on the 16 April 2020 (3407 sequences in total). Sequences were then aligned and sequences less than 10% divergent at the nucleotide level, identified using the ‘identify similar/ identical sequences’ function in SSE v1.4 were removed from the dataset. Sequences were annotated into animal groups and genera based on their description in the NCBI database. The trimmed dataset (**Table S1**) included 215 complete genome coronavirus sequences. Individual groups were made for sequences originating from the following hosts: bat (*n* = 108), avian (35), camelid (3), canine (7), feline (9), human (7), mustelids (5), rodents (8), swine (15), ungulates (9) and ‘other’ (which included bottle-nosed dolphin (2), hedgehog (2), rabbit (2), beluga whale (1), civet (1) and pangolin (1)). Groups were loosely defined based on taxonomic orders, with some exceptions made to examine our specific research questions. Bats are of the order Chiroptera; multiple avian orders were grouped together (Galliformes, Anseriformes, Passeriformes, Gruiformes, Columbiformes and Pelicaniformes); even toed (Artiodactyla) and odd toed (Perissodactyla) ungulate orders were grouped, with camelids analysed separately due to their association with MERS-CoV (Azhar, et al. 2014); Canidae (canine) and Pantherinae (feline) sequences of the Carnivora order were analysed separately, as canines have previously been suggested as an intermediate host species for SARS-CoV-2 (Xia 2020) and cat infections with SARS-CoV-2 have been reported (Shi, et al. 2020); humans were the only representatives from the Primate order; all remaining Carnivora, with the exception of a single civet sequence, belonged to the Mustelidae (mustelids); rodents belong to the Rodentia order; and swine belong to the Artiodactyla order; whales are also Artodactyla but swine were considered separately due to considerable interest in porcine coronaviruses (Vlasova, et al. 2020). Sequences were also annotated for genus by reference to the NCBI description (203 of the 215 sequences were assigned to a genus), and for primary replication site by literature reference (refer to **Table S1**). Replication site annotations were based on the sample type from which a coronavirus sequence was obtained – ‘enteric’ for faecal/ gastrointestinal samples, ‘respiratory’ for nasal, oropharyngeal and other respiratory samples; ‘multiple’ if samples from multiple systems tested positive, ‘other’ if the sample was collected from a site not falling into the enteric or respiratory categories (e.g. brain), or ‘unknown’ if a sample type could not be determined. If only one sampling route was tested and returned a positive result, the sequence was categorised in accordance with the sole sampling route. The sequence datasets used in this paper are summarised in **Fig. 1**.

### Analyses of dinucleotide content

CpG and UpA composition of complete genomes or of individual ORFs were calculated using the composition scan in SSE v1.4. CpG frequencies were measured as observed: expected (O:E) ratios, using the formula f(CpG)/ f(C)*f(G). Individual ORFs were identified using a combination of ORF finder (https://www.ncbi.nlm.nih.gov/orffinder/), visual inspection of nucleotide alignments in SSE v1.4, comparison with previous literature and information available from nextstrain.org. Sliding window analyses were performed on the 1163 aligned SARS-CoV-2 sequences and the related bat and pangolin sequences by performing composition scans in SSE v1.4 for 100 nucleotide genomic regions, at 25 nucleotide iterations. For the SARS-CoV-2 sequences, mean CpG O:E ratios for each window were calculated. CPB (Gutman and Hatfield 1989) scores across the SARS-CoV-2 ORFeome were calculated using the SSE v1.4 composition scan function. Individual ORFs were concatenated with a separating ‘NNN’ codon for analysis, and secondary overlapping ORFs were not included due to coding constraints imposed in these regions.

### Phylogenetic analyses

E ORFs were aligned in MEGA X (Kumar, et al. 2018) using the Clustal method. Of the 215 divergent sequences included in the analysis, E ORF could be identified in 178 by homology with E ORFs previously annotated in NCBI. Of these 178 E ORFs, 7 were sequences isolated from humans and 96 were from bats; these sequences were selected for analysis. Phylogenetic reconstruction was performed using an unrooted maximum likelihood tree, with gamma distributed variation in rates between branches and 100 bootstraps (also in MEGA X).

### Statistical analyses

Comparison to determine whether there was a statistically significant difference across groups was performed using a 1-way ANOVA in GraphPad Prism. Multi-way comparisons of CpG composition between groups were performed using Tukey’s multiple comparisons test in GraphPad Prism. A 5% significance level was used throughout.

## Acknowledgements

We would like to thank Prof. Peter Simmonds (Oxford University) for providing a pre-release version of SSE v1.4. We are grateful to Dr James Glover (the Roslin Institute) for comments on the manuscript. Figure 1 was created using BioRender. This work was supported by BBSRC Roslin Institute Strategic Program Grant funding (no. BB/P013740/1 to PD and FG) and Wellcome Trust/ Royal Society Fellowship (211222/Z/18/Z to EG).

## References

Atkinson N, Witteveldt J, Evans D, Simmonds P. 2014. The influence of CpG and UpA dinucleotide frequencies on RNA virus replication and characterization of the innate cellular pathways underlying virus attenuation and enhanced replication. Nucleic Acids Research 42:4527–4545.

Azhar EI, El-Kafrawy SA, Farraj SA, Hassan AM, Al-Saeed MS, Hashem AM, Madani TA. 2014. Evidence for Camel-to-Human Transmission of MERS Coronavirus. 370:2499–2505.

Banerjee A, Kulcsar K, Misra V, Frieman M, Mossman K. 2019. Bats and Coronaviruses. Viruses 11:41.

Berkhout B, van Hemert F. 2015. On the biased nucleotide composition of the human coronavirus RNA genome. Virus Research 202:41–47.

Burns CC, Campagnoli R, Shaw J, Vincent A, Jorba J, Kew O. 2009. Genetic Inactivation of Poliovirus Infectivity by Increasing the Frequencies of CpG and UpA Dinucleotides within and across Synonymous Capsid Region Codons. 83:9957–9969.

Chenik M, Chebli K, Blondel D. 1995. Translation initiation at alternate in-frame AUG codons in the rabies virus phosphoprotein mRNA is mediated by a ribosomal leaky scanning mechanism. 69:707–712.

Cooper DN, Krawczak M. 1989. Cytosine methylation and the fate of CpG dinucleotides in vertebrate genomes. Human Genetics 83:181–188.

Cotten M, Lam TT, Watson SJ, Palser AL, Petrova V, Grant P, Pybus OG, Rambaut A, Guan Y, Pillay D, et al. 2013. Full-genome deep sequencing and phylogenetic analysis of novel human betacoronavirus. Emerging infectious diseases 19:736–742B.

Cox E, Hooyberghs J, Pensaert MB. 1990. Sites of replication of a porcine respiratory coronavirus related to transmissible gastroenteritis virus. Research in veterinary science 48:165–169.

Davidson A, Williamson MK, Lewis S, Shoemark D, Carroll M, Heesom K, Zambon M, Ellis J, Lewis P, Hiscox J, et al. 2020. Characterisation of the transcriptome and proteome of SARS-CoV-2 using direct RNA sequencing and tandem mass spectrometry reveals evidence for a cell passage induced in-frame deletion in the spike glycoprotein that removes the furin-like cleavage site. In: bioRxiv.

Deaton AM, Bird A. 2011. CpG islands and the regulation of transcription. 25:1010–1022.

Dev RR, Ganji R, Singh SP, Mahalingam S, Banerjee S, Khosla S. 2017. Cytosine methylation by DNMT2 facilitates stability and survival of HIV-1 RNA in the host cell during infection. Biochemical Journal 474:2009–2026.

Ficarelli M, Wilson H, Pedro Galão R, Mazzon M, Antzin-Anduetza I, Marsh M, Neil SJD, Swanson CM. 2019. KHNYN is essential for the zinc finger antiviral protein (ZAP) to restrict HIV-1 containing clustered CpG dinucleotides. eLife 8:e46767.

Firth AE, Atkins JF. 2010. Candidates in Astroviruses, Seadornaviruses, Cytorhabdoviruses and Coronaviruses for +1 frame overlapping genes accessed by leaky scanning. Virology Journal 7:17.

Futcher B, Gorbatsevych O, Shen SH, Stauft CB, Song Y, Wang B, Leatherwood J, Gardin J, Yurovsky A, Mueller S, et al. 2015. Reply to Simmonds et al.: Codon pair and dinucleotide bias have not been functionally distinguished. 112:E3635–E3636.

Gaunt E, Wise HM, Zhang H, Lee LN, Atkinson NJ, Nicol MQ, Highton AJ, Klenerman P, Beard PM, Dutia BM, et al. 2016. Elevation of CpG frequencies in influenza A genome attenuates pathogenicity but enhances host response to infection. eLife 5:e12735–e12735.

Greenbaum BD, Levine AJ, Bhanot G, Rabadan R. 2008. Patterns of evolution and host gene mimicry in influenza and other RNA viruses. PLOS Pathogens 4:e1000079–e1000079.

Groenke N, Trimpert J, Merz S, Conradie AM, Wyler E, Zhang H, Hazapis O-G, Rausch S, Landthaler M, Osterrieder N, et al. 2020. Mechanism of Virus Attenuation by Codon Pair Deoptimization. Cell Reports 31:107586.

Guo X, Ma J, Sun J, Gao G. 2007. The zinc-finger antiviral protein recruits the RNA processing exosome to degrade the target mRNA. 104:151–156.

Gutman GA, Hatfield GW. 1989. Nonrandom utilization of codon pairs in Escherichia coli. 86:3699–3703.

Hiscott J, Kwon H, Génin P. 2001. Hostile takeovers: viral appropriation of the NF-kB pathway. The Journal of Clinical Investigation 107:143–151.

Hu B, Ge X, Wang L-F, Shi Z. 2015. Bat origin of human coronaviruses. Virology Journal 12:221.

Irigoyen N, Firth AE, Jones JD, Chung BYW, Siddell SG, Brierley I. 2016. High-Resolution Analysis of Coronavirus Gene Expression by RNA Sequencing and Ribosome Profiling. PLOS Pathogens 12:e1005473–e1005473.

Kanaya S, Yamada Y, Kinouchi M, Kudo Y, Ikemura T. 2001. Codon Usage and tRNA Genes in Eukaryotes: Correlation of Codon Usage Diversity with Translation Efficiency and with CG-Dinucleotide Usage as Assessed by Multivariate Analysis. Journal of Molecular Evolution 53:290–298.

Khoddami V, Cairns BR. 2013. Identification of direct targets and modified bases of RNA cytosine methyltransferases. Nature Biotechnology 31:458–464.

Kim D, Lee J-Y, Yang J-S, Kim JW, Kim VN, Chang H. 2020. The architecture of SARS-CoV-2 transcriptome.2020.2003.2012.988865.

Kozak M. 1986. Point mutations define a sequence flanking the AUG initiator codon that modulates translation by eukaryotic ribosomes. Cell 44:283–292.

Kumar S, Stecher G, Li M, Knyaz C, Tamura K. 2018. MEGA X: Molecular Evolutionary Genetics Analysis across Computing Platforms. Molecular Biology and Evolution 35:1547–1549.

Kunec D, Osterrieder N. 2016. Codon Pair Bias Is a Direct Consequence of Dinucleotide Bias. Cell Reports 14:55–67.

Liao CL, Lai MM. 1994. Requirement of the 5’-end genomic sequence as an upstream cis-acting element for coronavirus subgenomic mRNA transcription. 68:4727–4737.

Lin C-G, Lo SJ. 1992. Evidence for involvement of a ribosomal leaky scanning mechanism in the translation of the hepatitis B virus Pol gene from the viral pregenome RNA. Virology 188:342–352.

Lin Y-T, Chiweshe S, McCormick D, Raper A, Wickenhagen A, DeFillipis V, Gaunt E, Simmonds P, Wilson SJ, Grey F. 2020. Human cytomegalovirus evades ZAP detection by suppressing CpG dinucleotides in the major immediate early genes.2020.2001.2007.897132.

Lu R, Zhao X, Li J, Niu P, Yang B, Wu H, Wang W, Song H, Huang B, Zhu N, et al. 2020. Genomic characterisation and epidemiology of 2019 novel coronavirus: implications for virus origins and receptor binding. The Lancet 395:565–574.

Marra MA, Jones SJM, Astell CR, Holt RA, Brooks-Wilson A, Butterfield YSN, Khattra J, Asano JK, Barber SA, Chan SY, et al. 2003. The Genome Sequence of the SARS-Associated Coronavirus. 300:1399–1404.

McClelland M, Ivarie R. 1982. Asymmetrical distribution of CpG in an ‘average’ mammalian gene. Nucleic Acids Research 10:7865–7877.

Medvedeva YA, Khamis AM, Kulakovskiy IV, Ba-Alawi W, Bhuyan MSI, Kawaji H, Lassmann T, Harbers M, Forrest ARR, Bajic VB, et al. 2014. Effects of cytosine methylation on transcription factor binding sites. BMC Genomics 15:119.

Moratorio G, Henningsson R, Barbezange C, Carrau L, Bordería AV, Blanc H, Beaucourt S, Poirier EZ, Vallet T, Boussier J, et al. 2017. Attenuation of RNA viruses by redirecting their evolution in sequence space. Nature Microbiology 2:17088.

O’Connor JB, Brian DA. 2000. Downstream Ribosomal Entry for Translation of Coronavirus TGEV Gene 3b. Virology 269:172–182.

Odon V, Fros JJ, Goonawardane N, Dietrich I, Ibrahim A, Alshaikhahmed K, Nguyen D, Simmonds P. 2019. The role of ZAP and OAS3/RNAseL pathways in the attenuation of an RNA virus with elevated frequencies of CpG and UpA dinucleotides. Nucleic Acids Research 47:8061–8083.

Perlman S, Netland J. 2009. Coronaviruses post-SARS: update on replication and pathogenesis. Nature Reviews Microbiology 7:439–450.

Rasschaert D, Duarte M, Laude H. 1990. Porcine respiratory coronavirus differs from transmissible gastroenteritis virus by a few genomic deletions. 71:2599–2607.

Rima BK, McFerran NV. 1997. Dinucleotide and stop codon frequencies in single-stranded RNA viruses. 78:2859–2870.

Rota PA, Oberste MS, Monroe SS, Nix WA, Campagnoli R, Icenogle JP, Peñaranda S, Bankamp B, Maher K, Chen M-h, et al. 2003. Characterization of a Novel Coronavirus Associated with Severe Acute Respiratory Syndrome. 300:1394–1399.

Ryabova LA, Pooggin MM, Hohn T. 2006. Translation reinitiation and leaky scanning in plant viruses. Virus Research 119:52–62.

Sawicki SG, Sawicki DL. 1998. A New Model for Coronavirus Transcription. In: Enjuanes L, Siddell SG, Spaan W, editors. Coronaviruses and Arteriviruses. Boston, MA: Springer US. p. 215–219.

Sawicki SG, Sawicki DL, Siddell SG. 2007. A Contemporary View of Coronavirus Transcription. 81:20–29.

Schaecher SR, Mackenzie JM, Pekosz A. 2007. The ORF7b Protein of Severe Acute Respiratory Syndrome Coronavirus (SARS-CoV) Is Expressed in Virus-Infected Cells and Incorporated into SARS-CoV Particles. 81:718–731.

Schneider PA, Kim R, Lipkin WI. 1997. Evidence for translation of the Borna disease virus G protein by leaky ribosomal scanning and ribosomal reinitiation. 71:5614–5619.

Senanayake SD, Brian DA. 1997. Bovine coronavirus I protein synthesis follows ribosomal scanning on the bicistronic N mRNA. Virus Research 48:101–105.

Shi J, Wen Z, Zhong G, Yang H, Wang C, Huang B, Liu R, He X, Shuai L, Sun Z, et al. 2020. Susceptibility of ferrets, cats, dogs, and other domesticated animals to SARS–coronavirus 2.eabb7015.

Simmonds P. 2012. SSE: a nucleotide and amino acid sequence analysis platform. BMC Research Notes 5:50.

Simmonds P, Xia W, Baillie JK, McKinnon K. 2013. Modelling mutational and selection pressures on dinucleotides in eukaryotic phyla –selection against CpG and UpA in cytoplasmically expressed RNA and in RNA viruses. BMC Genomics 14:610.

Squires JE, Patel HR, Nousch M, Sibbritt T, Humphreys DT, Parker BJ, Suter CM, Preiss T. 2012. Widespread occurrence of 5-methylcytosine in human coding and non-coding RNA. Nucleic Acids Research 40:5023–5033.

Sved J, Bird A. 1990. The expected equilibrium of the CpG dinucleotide in vertebrate genomes under a mutation model. 87:4692–4696.

Takata MA, Gonçalves-Carneiro D, Zang TM, Soll SJ, York A, Blanco-Melo D, Bieniasz PD. 2017. CG dinucleotide suppression enables antiviral defence targeting non-self RNA. Nature 550:124–127.

Tang Q, Wang X, Gao G. 2017. The Short Form of the Zinc Finger Antiviral Protein Inhibits Influenza A Virus Protein Expression and Is Antagonized by the Virus-Encoded NS1. 91:e01909–01916.

Tats A, Tenson T, Remm M. 2008. Preferred and avoided codon pairs in three domains of life. BMC Genomics 9:463.

Tomso DJ, Bell DA. 2003. Sequence Context at Human Single Nucleotide Polymorphisms: Overrepresentation of CpG Dinucleotide at Polymorphic Sites and Suppression of Variation in CpG Islands. Journal of Molecular Biology 327:303–308.

Tulloch F, Atkinson NJ, Evans DJ, Ryan MD, Simmonds P. 2014. RNA virus attenuation by codon pair deoptimisation is an artefact of increases in CpG/UpA dinucleotide frequencies. eLife 3:e04531.

van der Hoek L, Pyrc K, Jebbink MF, Vermeulen-Oost W, Berkhout RJM, Wolthers KC, Wertheim-van Dillen PME, Kaandorp J, Spaargaren J, Berkhout B. 2004. Identification of a new human coronavirus. Nature Medicine 10:368–373.

Vlasova AN, Wang Q, Jung K, Langel SN, Malik YS, Saif LJ. 2020. Porcine Coronaviruses. Emerging and Transboundary Animal Viruses:79–110.

Wise HM, Barbezange C, Jagger BW, Dalton RM, Gog JR, Curran MD, Taubenberger JK, Anderson EC, Digard P. 2011. Overlapping signals for translational regulation and packaging of influenza A virus segment 2. Nucleic Acids Research 39:7775–7790.

Woo PCY, Lau SKP, Chu C-m, Chan K-h, Tsoi H-w, Huang Y, Wong BHL, Poon RWS, Cai JJ, Luk W-k, et al. 2005. Characterization and Complete Genome Sequence of a Novel Coronavirus, Coronavirus HKU1, from Patients with Pneumonia. 79:884–895.

Woo PCY, Wong BHL, Huang Y, Lau SKP, Yuen K-Y. 2007. Cytosine deamination and selection of CpG suppressed clones are the two major independent biological forces that shape codon usage bias in coronaviruses. Virology 369:431–442.

Xia X. 2020. Extreme genomic CpG deficiency in SARS-CoV-2 and evasion of host antiviral defense. Molecular Biology and Evolution.

Xie L, Lu B, Zheng Z, Miao Y, Liu Y, Zhang Y, Zheng C, Ke X, Hu Q, Wang H. 2018. The 3C protease of enterovirus A71 counteracts the activity of host zinc-finger antiviral protein (ZAP). 99:73–85.

Zaki AM, van Boheemen S, Bestebroer TM, Osterhaus ADME, Fouchier RAM. 2012. Isolation of a Novel Coronavirus from a Man with Pneumonia in Saudi Arabia. 367:1814–1820.

Zhu N, Zhang D, Wang W, Li X, Yang B, Song J, Zhao X, Huang B, Shi W, Lu R, et al. 2020. A Novel Coronavirus from Patients with Pneumonia in China, 2019. New England Journal of Medicine 382:727–733.

